# Development of a 16S metabarcoding assay for the environmental DNA (eDNA) detection of aquatic reptiles across northern Australia

**DOI:** 10.1101/2020.09.29.319525

**Authors:** Katrina West, Matthew Heydenrych, Rose Lines, Tony Tucker, Sabrina Fossette, Scott Whiting, Michael Bunce

## Abstract

A severe lack of distribution data for aquatic reptiles in northern Australia leaves many taxa vulnerable to extirpation and extinction. Environmental DNA (eDNA) technologies offer sensitive and non-invasive genetic alternatives to trapping and visual surveys and are increasingly employed for the detection of aquatic and semi-aquatic reptiles. However, at present, these studies have largely applied species-specific primers which do not provide a cost-effective avenue for the simultaneous detection of multiple reptilian taxa. Here, we present a 16S rRNA metabarcoding assay for the broad detection of aquatic and semi-aquatic reptile species. This assay is tested on water samples collected at multiple sampling sites at two tropical locations: 12 marine/estuarine sites in Roebuck Bay, Western Australia, and 4 estuarine sites in Cooktown, Queensland, Australia. A total of nine reptile taxa were detected from 10 of the 16 sampled sites, including marine and freshwater turtles, aquatic and semi-aquatic/terrestrial snakes, and terrestrial skinks. However, inconsistencies in the detection of previously observed aquatic reptiles at our sampled sites, such as saltwater crocodile and sea snakes, indicates that further research is required to assess the reliability, strengths and limitations of eDNA methods for aquatic reptile detection before it can be integrated as a broad-scale bioassessment tool.

## 1. Introduction

The aquatic and semi-aquatic reptile fauna of northern Australia are highly distinctive, exhibiting life history traits and physical adaptations to extreme climates and, in the case of semi-aquatic species, seasonal, but increasingly inconsistent, water availability (Pusey, 2011). Collectively these taxa form an integral component of the region’s aquatic and riparian food webs across multiple trophic levels; the saltwater crocodile (*Crocodylus porosus*) often fills the role of apex predator in these ecosystems. There are over 90 recognised aquatic and semi-aquatic reptile species in Australia, including marine and freshwater turtles, crocodiles, monitor lizards, water skinks, water dragons, sea snakes, sea kraits and other semi-aquatic snakes (Uetz, Freed and Hošek, 2018). Whilst their terrestrial counterparts have not historically experienced population declines in Australia – largely reflecting their inhabitance of arid, inland areas less disturbed by anthropogenic influences (Fox, 2008), aquatic and semi-aquatic reptiles species are under greater threat from pollution, urban development, over-harvesting, fisheries bycatch, invasive species (such as the toxic cane toad *Rhinella marina*), and climate change; the latter affecting the frequency of bushfires, coastal erosion and coral reef degradation (Böhm et al., 2013; Doody et al., 2014; Milton, 2001; Wilcox et al., 2015). In addition, latitudinal niche shifts to mitigate changing temperatures have prehistorically been much more challenging for ectotherms, such as reptiles, which have a lower climatic tolerance and reproductive ability (Rolland *et al.*, 2018). The viability of reptile species that exhibit temperature-dependent sex determination may be compromised with projected temperature shifts in the next 50 years (Tomillo *et al.*, 2015). Thus, the continual monitoring of aquatic and semi-aquatic reptile species, particularly in northern Australia, is an increasing necessity.

Typical survey techniques for the detection of aquatic reptiles include snorkelling, trapping, satellite tracking, aerial surveying, seining and bycatch reports, although many of these invasive methods can be limited in northern Australian waterbodies by the threat posed by saltwater crocodiles (Australian Government, 2011). In comparison to other faunal groups, there is a severe lack of data on aquatic reptile distributions across catchments in northern Australia (Fox, 2008). The majority of aquatic reptile surveys in this region have targeted the saltwater crocodile; two thirds of consolidated aquatic/semi-aquatic reptile distribution records are attributed to this species (Fox, 2008), leaving data deficiencies for other reptilian taxa.

The coastal and offshore waters of northern Australia are a biodiversity hotspot for sea snakes (Elapidae: Hydrophiinae) with at least 32 species, of which 4 are nationally endemic (Uetz, Freed and Hošek, 2018). Two of these endemic species, *Aipysurus foliosquama* and *A. apraefrontalis*, were elevated to ‘Critically Endangered’ in 2011 under Australia’s Environment Protection and Biodiversity Conservation (EPBC) Act 1999, given sustained declines on the Ashmore and Hibernia Reefs, with a total estimated area of occupancy totalling only 227 km^2^ (Guinea, 2003; Guinea & Whiting, 2005; Lukoschek, Beger, Ceccarelli, Richards, & Pratchett, 2013). There is a lack of robust distribution data for sea snakes globally; 34% of all sea snakes are listed as Data Deficient (DD) on the IUCN Red List (Elfes *et al.*, 2013). Data deficiencies for sea snakes in northern Australia include the Arafura sea snake (*Aipysurus tenuis*), Zweifel’s sea snake (*Enhydrina zweifeli*), the faint-banded sea snake (*Hydrophis belcheri*), the fine-spined sea snake (*Hydrophis czeblukovi*), the slender-necked sea snake (*Hydrophis melanocephalus*) and the northern mangrove sea snake (*Parahydrophis mertoni*; Elfes et al., 2013). Other DD aquatic reptile taxa in northern Australia include freshwater turtles, such as Irwin’s snapping turtle (*Elseya irwini*), the Gulf snapping turtle (*Elseya lavarackorum*), and the northern yellow-faced turtle (*Emydura tanybaraga*; Van Dyke, Ferronato, & Spencer, 2018), and the marine flatback turtle (*Natator depressus*).

Whilst a high level of data deficiency does not necessarily correspond directly to elevated extinction risk, insufficient information in regards to population trajectories, distribution and taxonomy creates a lot of uncertainty around extinction risk, conservation priorities and legislation (Böhm *et al.*, 2013; Bland and Böhm, 2016). Robust temporal and spatial distribution records provide the baseline upon which species ecology and conservation status can (and must) be developed. However, given the sheer number of DD reptile taxa, that may or may not be threatened, it remains economically challenging to conduct in-depth surveys. Furthermore, DD taxa may require more specialised trapping techniques, taxonomic expertise and may react poorly to trapping/handling. In addition, rare and cryptic taxa may never be detected in timeframes that are required for management decisions, such as in relation to coastal development assessments.

Environmental DNA (eDNA) technologies offer a sensitive, cost-effective and non-invasive genetic alternative to individual species and multi-taxon surveying in marine, freshwater and terrestrial environments (Thomsen *et al.*, 2012; Bohmann *et al.*, 2014; Evans *et al.*, 2016; Olds *et al.*, 2016). To date, reptile eDNA studies have largely applied species-specific markers to amplify individual species from mixed environmental samples (Davy, Kidd and Wilson, 2015; De Souza *et al.*, 2016; Halstead *et al.*, 2017; Baker *et al.*, 2018; Feist *et al.*, 2018; Ratsch, Kingsbury and Jordan, 2020; Rose, Fukuda and Campbell, 2020). The first reptile eDNA study, published in 2014, developed a diagnostic PCR to detect the Burmese python (*Python bivittatus*), a semi-aquatic, invasive species in Florida (Piaggio *et al.*, 2014). The python’s elusive nature, cryptic colouration and occupation of aquatic habitats that were logistically difficult to survey, prompted an eDNA approach. Piaggio et al., (2014) developed a *P. bivittatus*-specific mitochondrial cytochrome b assay that was applied to water samples from field sites in south Florida, successfully detecting the species where it had been previously observed.

Advances in high-throughput sequencing now allow the simultaneous amplification and sequencing of multiple taxa through universal or broad-taxonomic PCR assays (referred to as eDNA metabarcoding), proving a more efficient approach to genetic surveying. However, the use of eDNA metabarcoding for the detection of reptile assemblages has not yet been thoroughly tested. Kelly, Port, Yamahara, & Crowder (2014) applied vertebrate-specific mitochondrial 12S rRNA primers to detect green sea turtle (*Chelonia mydas*) in a large mesocosm, but were unsuccessful, despite successfully amplifying the intended target with species-specific primers. Conversely, Lacoursière-Roussel, Dubois, Normandeau, & Bernatchez (2016) had more success in using COI metabarcoding assays to detect three species of snake and two species of turtle across lakes and rivers in Canada. Our primary objectives in this study were to design a metabarcoding assay that is able to simultaneously target aquatic and semi-aquatic reptile groups in northern Australia and test its utility on water samples collected across northern Australia.

## 2. Materials and methods

### 2.1 Field sampling

A total of 22 one-litre surface water samples were collected at 12 sites in Roebuck Bay, Western Australia in August 2018 and 20 one-litre surface water samples collected at four sites near Cooktown, Queensland in March/April 2020 (Figure 1; Table S1). Roebuck Bay is a semiarid, tropical, marine embayment characterised by intertidal sand, mudflat and mangrove habitats. Surface water samples were collected during the dry season at eight ocean sites, two estuarine creek sites, and two intertidal/mangrove sites. There were a number of marine turtle and sea snake species observed during sampling across the majority of sites, whilst saltwater crocodiles have been previously observed in the vicinity of the two sampled creek sites (Table S1). The Cooktown region is comprised of a variety of tropical landscapes such as sandy beaches, tidal estuaries, freshwater wetlands and rainforest hinterland. Surface water samples were collected in the wet season at four estuarine creek sites that outflow into the ocean; two of which contained mangrove vegetation and two that were connected to a sandy wetland system. The Cooktown surface water samples were collected using a large pole to mitigate saltwater crocodile risks in the area. Water samples were individually filtered across Pall 0.45μm GN-6 Metricel® mixed cellulose ester membranes using a Pall Sentino® Microbiology pump (Pall Corporation, Port Washington, USA). Filter membranes were immediately frozen and stored at -20°C prior and post-transportation to the Trace & Environmental DNA (TrEnD) Laboratory in Perth, Western Australia.

**Figure 1.**
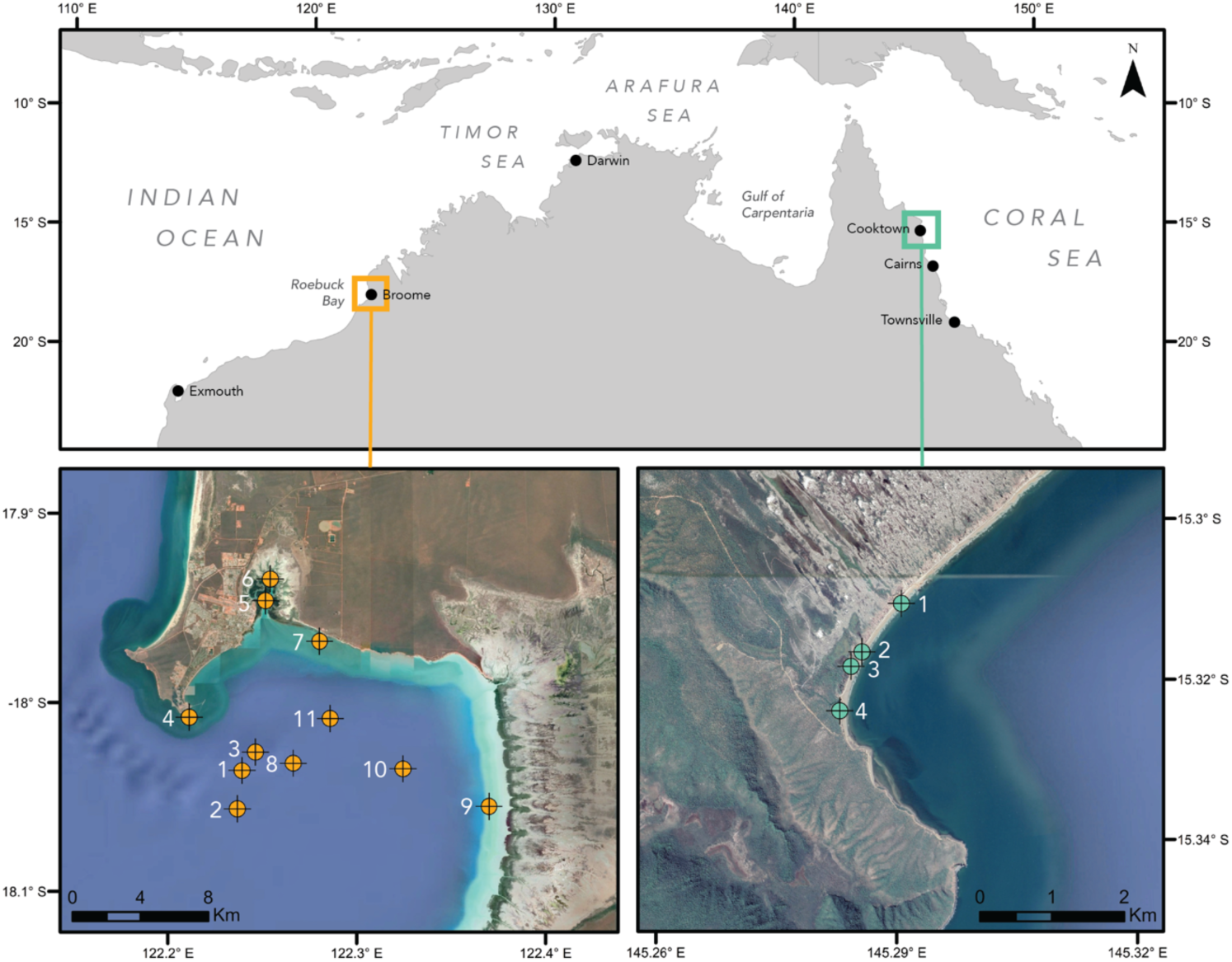
Location of sampling sites across northern Australia. One-litre water samples were collected at 12 sites in Roebuck Bay, Western Australia (n=22; bottom-left map) and from 4 sites near Cooktown, Queensland (n=20; bottom-right map). Further site information is provided in Table S1. Map data: Google Earth, SIO, NOAA, U.S. Navy, NGA, GEBCO; image: Landsat/Copernicus (bottom-left map), CNES/Airbus (bottom-right map).

### 2.2 *In-silico* design

Mitochondrial DNA is typically targeted for eDNA metabarcoding assays due to both template copy number and reference data. Two Indo-Pacific sea snake databases were curated for the mitochondrial 16S rDNA and cytochrome b gene regions by Sanders, Lee, Bertozzi, & Rasmussen (2013) and provided for this project. The cytochrome b database was substantially larger (361 sequences, 1104bp length) than the 16S database (50 sequences, 531bp length), however the cytochrome b region presented no conserved regions that were consistent across all sea snake sequences and that would be suitable for primer binding. Conversely, the 16S region exhibited a few conserved regions (approximately 30-50bp length) flanking larger hypervariable regions. Additional 16S rDNA sequences of a number of Australian aquatic and semi-aquatic reptiles (Table 1) were downloaded from NCBI GenBank (https://www.ncbi.nlm.nih.gov/genbank/) and aligned using the MUSCLE plugin in Geneious 10.0.6. Primer pairs were designed in conserved regions using the built-in primer design tool in Geneious; the final primer pair chosen by visually inspecting alignments for a target region with maximum variation and under a length of 280bp. The length constriction allows for the sequencing of barcode indexes (on the 5’ end of a forward/reverse primer within a fusion-tagged primer setup) on a single-end Illumina MiSeq sequencing run (up to 325bp). The forward primer was modified to include a degenerate base allowing for annealing to polymorphic sites in the reptile alignment. The resulting primer pair (herein referred to as the 16S Reptile assay) are AqReptileF-degenerate: 5’– AGACNAGAAGACCCTGTG-3’ and AqReptileR: 5’–CCTGATCCAACATCGAGG-3’; with a G/C content between 50.0-55.6%, T_M_ between 52.0-55.5 and hairpin T_M_ of 32.4.

**Table 1.** List of targeted Australian aquatic and semi-aquatic reptile species for in-silico primer analysis. ^ Indicates selection of reference sequences sourced from Sanders et al., (2013). * Indicates species whose tissue extracts were used to test primers *in vitro*. Target length refers to length of amplicon, ignoring primers. Under mismatches, F refers to forward primer mismatches.

| Common name | Species name | GenBank accession number | Target length (bp) | Mismatches |
| --- | --- | --- | --- | --- |
| **Sea snake** |  |  |  |  |
| ^Short-nosed sea snake | *Aipysurus apraefrontalis* | JX423420 | 213 | 0 |
| ^Dusky sea snake | *Aipysurus fuscus* | JX423430 | 213 | 0 |
| ^Olive sea snake | *Aipysurus laevis* | EU547181 | 213 | 0 |
| ^Turtle-headed sea snake | *Emydocephalus annulatus* | EU547185 | 214 | 0 |
| ^Black-headed sea snake | *Hydrophis atriceps* | KC014320 | 212 | 0 |
| ^Blue-banded sea snake | *Hydrophis cyanocinctus* | KC014331 | 212 | 0 |
| ^Ornate sea snake | *Hydrophis ornatus* | KC014358 | 212 | 0 |
| ^Yellow-bellied sea snake | *Pelamis platurus* | KC014375 | 212 | 0 |
| Stokes's sea snake | *Hydrophis stokesii* | JQ217146 | 212 | 0 |
| Horned sea snake | *Hydrophis peronii* | KU323976 | 212 | 0 |
| Shaw's sea snake | *Hydrophis curtus* | KX239662 | 212 | 0 |
| **Sea krait** |  |  |  |  |
| Yellow-lipped sea krait | *Laticauda colubrine* | NC_036054 | 212 | 0 |
| **Crocodile** |  |  |  |  |
| *Saltwater crocodile | *Crocodylus porosus* | NC_008143 | 275 | 0 |
| Freshwater crocodile | *Crocodylus johnstoni* | NC_015238 | 275 | 0 |
| **Marine turtle** |  |  |  |  |
| *Flatback turtle | *Natator depressus* | NC_018550 | 259 | 0 |
| Loggerhead turtle | *Caretta caretta* | MF579505 | 258 | 0 |
| Olive ridley turtle | *Lepidochelys olivacea* | DQ486893 | 258 | 0 |
| Leatherback turtle | *Dermochelys coriacea* | JX454992 | 259 | 0 |
| Green turtle | *Chelonia mydas* | JX454990 | 260 | 0 |
| Hawksbill turtle | *Eretmochelys imbricata* | MF571906 | 258 | 0 |
| **Freshwater turtle** |  |  |  |  |
| Pig-nosed turtle | *Carettochelys insculpta* | FJ862792 | 250 | 0 |
| Cann's snake-necked turtle | *Chelodina canni* | NC_041286 | 254 | 0 |
| Northern snake-necked turtle | *Chelodina rugosa* | KY776451 | 253 | 0 |
| Northern snapping turtle | *Elseya dentata* | KY779844 | 251 | 0 |
| Red-faced turtle | *Emydura victoriae* | NC_042473 | 253 | 0 |
| Western swamp turtle | *Pseudemydura umbrina* | NC_035731 | 244 | 0 |
| **Monitor lizard** |  |  |  |  |
| Mangrove monitor | *Varanus indicus* | EF193674 | 219 | 1 (F) |
| Argus monitor | *Varanus panoptes* | EF193685 | 223 | 1 (F) |

### 2.3 *In-vitro* testing on reptile tissue and eDNA water samples

*In-vitro* testing of the primers to assess PCR amplification and optimise the annealing temperature was firstly conducted using tissue extracts (1/10 dilution) from saltwater crocodile (*C. porosus*) and flatback turtle (*N. depressus*). The primers were then further tested on the filtered water samples collected at Roebuck Bay, Western Australia, and Cooktown, Queensland. DNA was extracted from half of the membrane using a DNeasy Blood and Tissue Kit (Qiagen; Venlo, Netherlands) with the following modifications: 540 μl of ATL lysis buffer, 60 μl of Proteinase K and a 3-hour digestion at 56°C. Blank controls were processed in parallel with all samples to detect any cross-contamination. Environmental DNA extracts were then stored at -20°C.

A gradient PCR determined an optimum annealing temperature of 52°C. Each qPCR reaction was carried out in 25μl containing: 1X AmpliTaq Gold® PCR buffer (Life Technologies, Massachusetts, USA), 2mM MgCl_2_, 0.1mM dNTPs, 0.2μM each of forward and reverse primers (Integrated DNA Technologies, Australia), 10ug BSA (Fisher Biotec, Australia), 0.6μl of 5X SYBR® Green (Life Technologies), 1U AmpliTaq Gold® DNA Polymerase (Life Technologies), 4μl of template DNA, and made to volume with Ultrapure™ Distilled Water (Life Technologies). Tissue and eDNA extracts were amplified in duplicate on a StepOnePlus Real-Time PCR System (Applied Biosystems, Massachusetts, USA) under the following conditions: initial denaturation at 95 °C for 5 min, followed by 50 cycles of 30 s at 95 °C, 52°C for 30 s and 45 s at 72 °C, with a final extension for 10 min at 72 °C. Quantitative PCR was performed in a single step using fusion tagged primer architecture that was comprised of a forward or reverse primer sequence, a unique index (6-8bp in length) and an Illumina compatible sequencing adaptor. All qPCR reactions were prepared in dedicated clean room facilities at the TrEnD Laboratory, Curtin University. Quantitative PCR amplicons were pooled at equimolar ratios based on their respective qPCR ΔRn values and were then size-selected (150-600bp) using a Pippin-Prep (Sage Science, Beverly, USA) to remove any off-target amplicons and primer dimer. Size-selected libraries were then purified using the Qiaquick PCR Purification Kit (Qiagen, Venlo, Netherlands), quantified using a Qubit 4.0 Fluorometer (Invitrogen, Carlsbad, USA) and diluted to 2 nM for loading onto a 300 cycle MiSeq® V2 Standard Flow Cell. Sequencing was conducted on an Illumina MiSeq platform (Illumina, San Diego, USA), housed in the TrEnD Laboratory at Curtin University, Western Australia.

Sequencing reads were demultiplexed using the ngsfilter (allowing up to three mismatches in primer sequences) and obisplit commands in the package OBITools (v1.2.9; Boyer et al., 2014) in RStudio (v1.1.423; RStudio Team, 2015). Data was then quality filtered (minimum length=100, maximum expected errors=2, no ambiguous nucleotides), denoised, filtered for chimeras and dereplicated (pool=TRUE) using the DADA2 bioinformatics package (Callahan et al., 2016) also implemented in RStudio. The resulting amplicon sequence variant (ASV) fasta file was queried against NCBI’s GenBank nucleotide database (accessed in 2020; Benson et al., 2005) using BLASTn (minimum percentage identity of 90, maximum target sequences of 10, reward value of 1) via Zeus, an SGI cluster, based at the Pawsey Supercomputing Centre in Kensington, Western Australia. Taxonomic assignments of ASVs were curated using a lowest common ancestor (LCA) approach (https://github.com/mahsa-mousavi/eDNAFlow/tree/master/LCA_taxonomyAssignment_scripts; Mousavi-Derazmahalleh et al., 2020, *in review*), whereby the top ten hits for each query are sequentially collapsed to the lowest common ancestor if the percentage identity between each consecutive hit differs by less than one percent (based on 100% query coverage). Each finalised taxonomic assignment therefore represents a query hit that is distinct from closely-related taxa.

## 3. Results

### 3.1 Primer design

The 16S Reptile assay was designed using sea snake, sea krait, crocodile, marine turtle, freshwater turtle and monitor lizard 16S rDNA reference sequences (Table 1). Target length of the amplified fragments ranged from 212bp (sea snake and sea krait) to 275bp (crocodile). There were no mismatches with the primers, except for the monitor lizard reference sequences (*Varanus indicus* and *Varanus panoptes*) which both exhibited one mismatch towards the 5’ end of the forward primer. It is unlikely that this will hinder potential amplification of monitor lizards with the 16S Reptile assay, but may impact on efficacy of detection when other (matching) reptile templates are more abundant.

The average pairwise percent identity for sea snakes across the target region was 94.2% (across 30 SNPs); the pairwise percent identity for congeneric sea snakes averaged 96.4% (min 93.4%, max 100%), providing enough variation to distinguish closely-related taxa. Across the two congeneric crocodiles and two congeneric monitor lizards, the pairwise percent identity was 95.3% (across 13 SNPs) and 88.8% (across 25 SNPs), respectively. For marine turtles (superfamily: Chelonioidea) the average pairwise percent identity was 90.9% (across 48 SNPs); and for freshwater turtles (family: Chelidae and Carettochelyidae) 74.0% (across 127 SNPs).

### 3.2 *In-vitro* primer testing

The 16S Reptile assay was firstly tested *in vitro* using tissue extractions (1/10 dilution) of salt water crocodile and flatback turtle. These successfully amplified with an average cycle threshold (C_T_) value of 27.1 and 21.9, respectively. The respective extracts matched to NCBI reference sequences of saltwater crocodile and flatback turtle with percentage identities of 100% and 99.6%.

Primers were then tested on the 46 water samples collected at Roebuck Bay and Cooktown. The majority of the eDNA extracts amplified with the 16S Reptile assay; albeit with high C_T_ values ranging from 28 to 39 reflecting low template copy numbers. The 16S Reptile assay yielded a total of 4,037,282 sequencing reads; the mean number of filtered sequences (post-quality, denoising and chimera filtering) was 17,796 ± 25,010 per sample. This resulted in a total of 96 taxa detected with the 16S Reptile assay (Table S2), with the majority of taxa being non-reptile (Figure 2). The highest average proportion of sequencing reads were attributed to bony fish (class: Actinopterygii, 36.7%), followed by amphibians (class: Amphibia, 33.6%), bivalve molluscs (class: Bivalvia, 12.9%), reptiles (class: Reptilia, 8.1%) and mammals (class: Mammalia, 8.1%). Two species of marine turtle (*N. depressus* and *Chelonia mydas*) were detected at 6/12 Roebuck Bay sites, whilst seven, largely freshwater-associated reptile taxa were detected at all four Cooktown creek sites (Table 2). The latter included freshwater turtles (*Myuchelys latisternum* and *Emydura*), aquatic and terrestrial snakes (Homalopsidae and *Dendrelaphis calligaster*) and skinks (*Saproscincus basiliscus, Carlia longipes* and *Carlia storri*).

**Table 2.** Reptile taxa detected from eDNA samples collected at Roebuck Bay (RB), Western Australia and Cooktown (CK), Queensland.

| Common name | Scientific name | Distribution | Site/s detected | Average read depth | Average % of total reads | Average % identity |
| --- | --- | --- | --- | --- | --- | --- |
| Indo-Australian water snakes | Homalopsidae | Northern Australia and South-east Asia. Widely located in both marine and freshwater, in addition to terrestrial environments. | CK 1 | 3247 | 24.2 | 94.6-94.8 |
| Northern tree snake | *Dendrelaphis calligaster* | Northern Australia, Indonesia, Papua New Guinea and Solomon Islands. Located in terrestrial environments with dense vegetation. | CK 3 | 92 | 1.0 | 95.8 |
| Pale-lipped shadenskink | *Saproscincus basiliscus* | Queensland, Australia. Located in rainforest environments. | CK 2 | 500 | 2.9 | 98.2 |
| Closed-litter rainbow-skink | *Carlia longipes* | Northern Queensland, Australia and southern Papua New Guinea. Located in open forest and rainforest environments. | CK 1 | 326 | 1.6 | 99.5 |
| Brown bicarinate rainbow-skink | *Carlia storri* | Located in Queensland and south-west Papua New Guinea. Located in the supralittoral zone, shrubland, savanna and forest environments. | CK 1 | 1773 | 9.1 | 98.1-98.6 |
| Flatback turtle | *Natator depressus* | Australian continental shelf. Located in soft-substrate and seagrass environments. Listed as 'Vulnerable' by the Australian EPBC Act 1999 and 'Data Deficient' by the IUCN. | RB 1, 3, 4, 5, 9, 11 | 198 | 11.6 | 99.6 |
| Green turtle | *Chelonia mydas* | Indo-Pacific and Atlantic. Located in hard and soft substrate, pelagic and seagrass environments. Listed as 'Vulnerable' by the Australian EPBC Act 1999 and 'Endangered' by the IUCN. | RB 4 | 16 | 0.5 | 100 |
| Saw-shelled turtle | *Myuchelys latisternum* | Northern and eastern Australia. Located in freshwater environments such as creeks, rivers, dams and lakes. | CK 1, 3, 4 | 166 | 1.6 | 100 |
| Australian short-necked turtles | *Emydura* | Northern and eastern Australia and in Papua New Guinea. Located in freshwater environments such as creeks, swamps, rivers, dams and lakes. | CK 4 | 126 | 2.6 | 97.2 |

**Figure 2.**
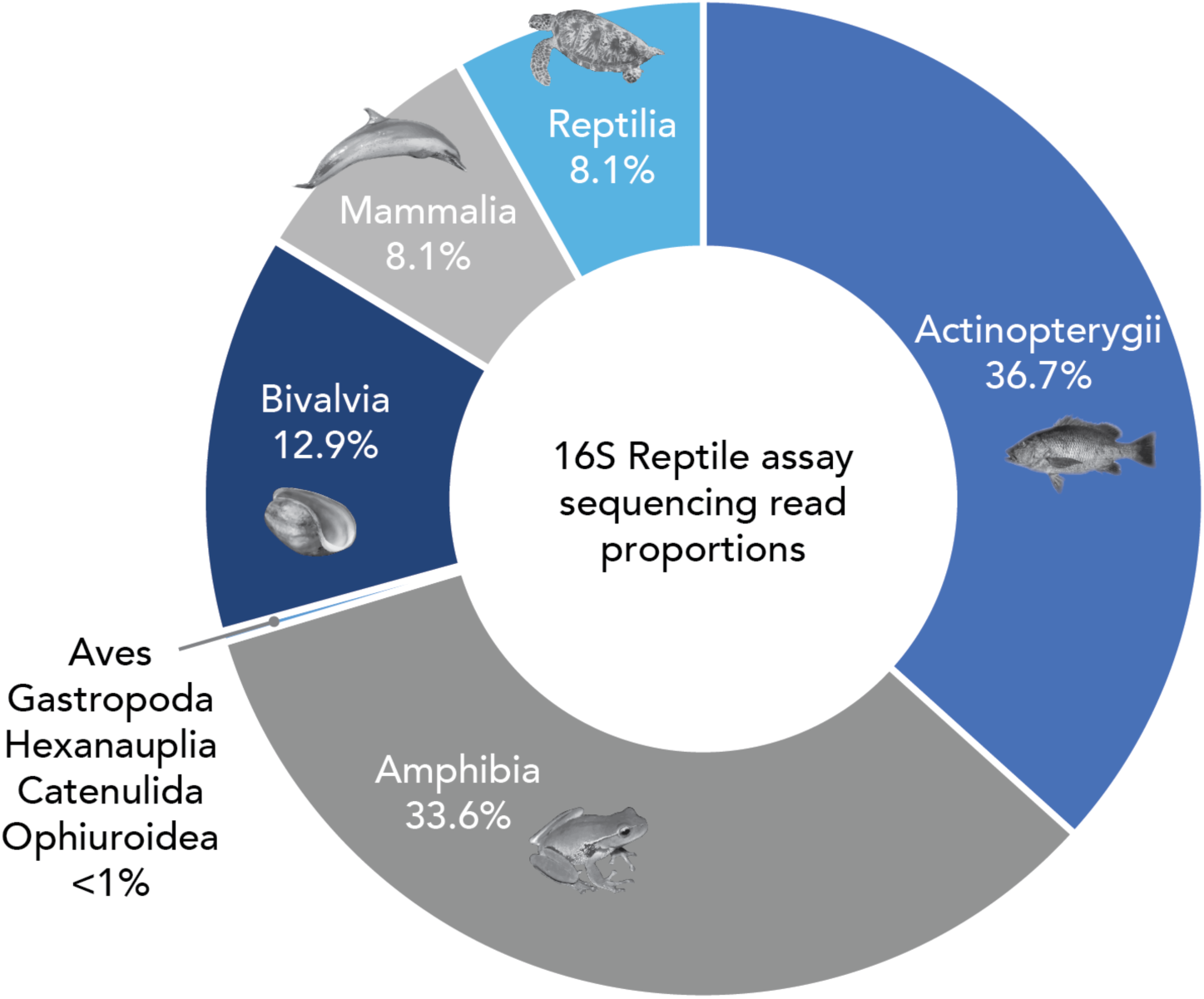
Proportion of Reptile 16S sequencing reads attributed at a class level from the eDNA study at Roebuck Bay, Western Australia and Cooktown, Queensland.

Average read depth of the detected reptiles varied from 16 reads (0.5% of average total reads) to 3247 reads (24.2% of average total reads). The percent identity match of the assigned reptile taxa ranged from 94.6% to 100% (Table 2). The majority of the species assignments had a high percent identity match (>98% with 100% query coverage). Only the northern tree snake (*D. calligaster*) had a lower percent identity match of 95.8% (with 100% query coverage). However, the queried eDNA sequence provided no other hits above 90%. This indicates that the eDNA sequence detected may represent intraspecific differentiation from the NCBI reference sequence of *D. calligaster*, or a closely-related taxon that is yet to be barcoded and shared on a publicly-accessible database. For both the genus and family assignments of *Emydura* and Homalopsidae respectively, there was not enough differentiation (≤1% percent identity difference) between closely-related taxa to confidently assign at a species level. However, for both queried eDNA sequences there were no percent identity matches above 98%, indicating that the detected sequences potentially represent taxa that are yet to be barcoded.

## 4. Discussion

The *in vitro* eDNA testing of our 16S Reptile metabarcoding assay successfully detected two marine turtle species at 6/12 Roebuck Bay sites, where they were visually observed in the area upon water sampling (Table S1). Flatback turtle (*Natator depressus*) was detected widely across ocean, creek and intertidal sites at Roebuck Bay, whilst green turtle (*Chelonia mydas*) was only detected at site 4 (ocean), despite being observed at creek and intertidal sites within Roebuck Bay. The principal detection of marine reptiles at Roebuck Bay was not unexpected, given that the surveyed sites were predominately marine-based with the exception of two estuarine sites. In comparison, the Cooktown sites, which were located solely within estuaries, provided a greater detection range of freshwater and also terrestrial reptile species. Two freshwater turtles (the saw-shelled turtle [*M. latisternum*] and an Australian short-necked turtle [genus: *Emydura*]) were detected at three of the Cooktown sites; these eDNA detections coincide within their known distribution ranges across northern and eastern Australia. Terrestrial skinks and a northern tree snake were additionally detected at the Cooktown sites, which likely reflects DNA shed into the sampled coastal creeks from drinking, skin shedding or other activities.

A notable detection at Cooktown was that of an Indo-Australian water snake (Homalopsidae). The family Homalopsidae is comprised of over 50 aquatic and semi-aquatic species that typically inhabit mangrove forests, tropical tidal wetlands and coastal waters from Southeast Asia to northern Australia (Alfaro *et al.*, 2008). The detected Homalopsidae eDNA sequence however could not be resolved to a species or even genus-level, as there were no high percentage matches (>98%) typically required for a species assignment, and of the remaining Homalopsidae hits (>90%) there was not enough resolution to confidently distinguish taxa even at the genus level. This indicates that the detected Homalopsidae eDNA sequence at Cooktown represents an Indo-Australian water snake that has not yet been barcoded for the 16S region and is potentially an undescribed species. Targeted herpetological surveying at this site is recommended to resolve this Homalopsidae eDNA detection.

A discrepancy in the performance of the 16S Reptile assay however, is that it did not detect any sea snake (Elapidae: Hydrophiinae) species, despite visual observations at 5 out of the 12 Roebuck Bay sites (Table S1). Additionally, it failed to detect any saltwater crocodile, despite previous observations in both the Roebuck Bay and Cooktown areas. This lack of detection could be attributed to the non-specificity of the assay and/or a low shedding rate of reptiles in the environment. Despite the fact that the assay was designed to preferentially amplify reptile taxa, reptiles only accounted for 8.1% of the average proportion of sequencing reads per sample, however this varied greatly between samples. The detection of the Indo-Australian water snake (Homalopsidae) for example, accounted for 24.2% of the total sequencing reads from Cooktown Site 1. Therefore, whilst the 16S Reptile assay does detect other taxonomic groups (bony fish, amphibians, bivalve molluscs and mammals), it is not exhibiting consistent preferential amplification of these groups above reptiles.

Further optimisation of this assay to reduce non-target amplification is ideal, although this may be complicated given that the primers are located in a highly-conserved region of the 16S rRNA gene. Furthermore, the inclusion of mismatches increases the risk that rare reptile variants will be excluded from amplification; additionally, the placement of mismatches on 3’ ends may compromise the efficiency of qPCR assays (Wilcox *et al.*, 2014). Alternatively, the development of discrete assays for each major reptilian order (i.e. turtles, crocodiles, tuatara, and squamates [lizards and snakes]) may be more effective in refining specificity, but retaining broad-taxonomic amplification. Another possibility is the use of blocking primers, which preferentially binds and restricts amplification of a targeted taxonomic group. Blocking primers have been successfully used in conjunction with metabarcoding assays to increase the specificity of amphibian and bony fish amplicons by reducing the amplification of human DNA (Valentini *et al.*, 2016; Sasso *et al.*, 2017).

The shedding rate of reptiles may also undermine our ability to detect them via eDNA methods. It has recently been suggested that reptiles may have a relatively lower shedding rate because of their keratinised scales and reduced urine production, and subsequently are less detectable than other mucus-covered organisms, such as fish and amphibians (Raemy and Ursenbacher, 2018; Adams *et al.*, 2019). Whilst this is yet to be explicitly tested, it may explain the inconsistent amplification of reptiles in aquatic environments across multiple studies. For example, giant garter snake (*Thamnophis gigas*) mesocosm experiments reported positive eDNA detection in tanks with snake skin and snake feces, however no detection with live snakes in tanks, nor at field locations, despite capture of the species with traps (Halstead *et al.*, 2017). The collection of water within one metre of eastern massasauga rattlesnakes (*Sistrurus catenatus*) only produced a positive eDNA detection for 2/100 water samples (Baker *et al.*, 2018). Conversely, Lacoursière-Roussel, Dubois, Normandeau, & Bernatchez (2016) successfully used metabarcoding to detect redbelly snake (*Storeria occipitomaculata*), northern watersnake (*Nerodia sipedon*), milksnake (*Lampropeltis triangulum*), snapping turtle (*Chelydra serpentine*) and wood turtle (*Glyptemys insculpta*) in rivers and lakes in Canada; however the wood turtle was not detected in 4 rivers that produced positive detections via species-specific qPCR and visual surveying. Overall, there has been a lot more success in the detection of wild turtles than snakes with eDNA, primarily with species-specific assays (Kelly *et al.*, 2014; De Souza *et al.*, 2016; Feist *et al.*, 2018), with a push to quantify turtle abundance and biomass using eDNA (Adams *et al.*, 2019). In regards to crocodiles, there is only one published study at present that has attempted to amplify crocodile eDNA. However despite observing West African crocodile (*Crocodylus suchus*) and the Nile monitor (*Varanus niloticus*) in the water at their field sites in Mauritania, they were unable to amplify any crocodile or other reptilian eDNA using a metabarcoding approach (Egeter *et al.*, 2018). A low shedding rate many therefore limit eDNA detection, despite a well-designed assay that is capable of amplifying the targeted taxa. It is possible that changing the eDNA substrate/method may help to improve detections by enriching fractions for target taxa – for example a plankton-tow might assist in retrieving reptile eDNA (Koziol et al., 2018). Nonetheless, an increase in sampling density may be the most feasible approach to increase eDNA sensitivity for aquatic reptiles.

Another limitation to the implementation of eDNA metabarcoding for broad-reptile surveying is potential reference database gaps for DD taxa. At present, only 27.6% of reptile species have been barcoded for the mitochondrial cytochrome *c* oxidase I (COI) gene; a short, standardised gene region that has historically formed the primary barcode sequence for animal species (Ratnasingham and Hebert, 2007). Only four sea snake species (Elapidae: Hydrophiinae) have been barcoded for this region, and only two of which are distributed in northern Australia. Our assay design targeted the mitochondrial 16S rDNA gene region, given the development of a 16S Indo-Pacific sea snake database with a high representation of northern Australian sea snakes (Sanders *et al.*, 2013). The development and implementation of eDNA metabarcoding for reptile species, particularly for DD taxa, should ideally be tailored to available reference sequences for the targeted taxonomic group. As such, the development of discrete taxonomic assays, as discussed previously for refining specificity, may be a superior approach to ensure high-resolution assignments (i.e. to a species-level). Ultimately however, it will be easier to implement broad metabarcoding assays with either a standardised barcode region, a complete suite of barcoded gene regions, or ideally, a complete mitochondrial genome for each representative species.

The inconsistent amplification and detection of aquatic reptilian eDNA, despite positive visual and trapping detections at survey sites, indicates that at present eDNA methodology provides an unreliable estimate of diversity and community composition between sites. We recommend that aquatic and semi-aquatic reptile shedding rates into various substrates (e.g. water, sediment, soil) are substantially tested before eDNA approaches, in particular metabarcoding, are further applied as reptile survey tools. This will provide greater insight into inconsistencies in amplification between taxonomic groups and whether assays need to be tailored to accommodate this i.e. the use of species-specific assays for taxa with low shedding rates. An alternative approach to detecting reptiles with potentially low shedding rates, would be to explore sampling volumes and subsequent filtering methods, i.e. increasing our standard water replicate volumes from 1L up to 50L. Here, we present a 16S rDNA primer assay for the broad detection of aquatic and semi-aquatic reptile species in northern Australia. However, constrictions around suitable primer binding regions that can simultaneously amplify deeply diverged reptile lineages has resulted in non-target amplification of other closely-related metazoan groups, such as amphibians. If a higher level of specificity is desired, we further recommend that reptile eDNA metabarcoding assays are developed at an order level or lower, and consider the coverage of reference databases for various gene regions. Taken together, we advocate that this 16S reptile assay is a valuable addition to the metabarcoding assay ‘toolkit’ and like many of the other assays developed (Miya *et al.*, 2015; Elbrecht and Leese, 2017; Taberlet *et al.*, 2018; Nester *et al.*, 2020) will be useful when designing or screening environmental samples for reptiles and other taxa.

## 5. Acknowledgements

No permits were necessary for water sampling but the Roebuck Bay study occurred within the Roebuck Bay Marine Park with the assistance of NBY (Nyamba Buru Yawuru) Country Managers and the WA Department of Biodiversity, Conservation and Attractions (DBCA) Marine Park rangers. Vessels and coxswains were supervised by the DBCA in Broome. We acknowledge eDNA frontiers and client for sourcing Cooktown water samples. For access to the Zeus supercomputer, which sped up much of our bioinformatic processing, we would like to thank Pawsey Supercomputing Centre (Kensington, WA). Lastly, we would like to give thanks to everyone at the TrEnD Laboratory for invaluable eDNA assistance across the duration of the project. This research did not receive any specific funding.

## 6. Conflicts of interest

The authors declare no conflicts of interest.

## Notes

### Competing Interest Statement

The authors have declared no competing interest.

