## Supplementary Information for "Development of a 16S metabarcoding assay for the environmental DNA (eDNA) detection of aquatic reptiles across northern Australia"

### Table S1. Site location information for eDNA water samples

| Location | Site # | Coordinates Latitude | Longitude | No. 1L samples | Habitat | Date Collected | Additional notes |
|---|---|---|---|---|---|---|---|
| Roebuck Bay | 1 | -18.0360 | 122.2392 | 2 | Ocean | 28/08/2018 | Sea snake and flatback turtle were observed |
| Roebuck Bay | 2 | -18.0564 | 122.2367 | 2 | Ocean | 28/08/2018 | Olive ridley turtle observed |
| Roebuck Bay | 3 | -18.0261 | 122.2462 | 2 | Ocean | 28/08/2018 | Sea snake and flatback turtle observed |
| Roebuck Bay | 4 | -18.0077 | 122.2113 | 1 | Ocean | 28/08/2018 | |
| Roebuck Bay | 5 | -17.9462 | 122.2516 | 2 | Creek | 29/08/2018 | Green turtle observed. Crocodiles have been previously observed here. |
| Roebuck Bay | 6 | -17.9347 | 122.2541 | 2 | Creek | 29/08/2018 | Green turtle observed. Crocodiles have been previously observed here. |
| Roebuck Bay | 7 | -17.9676 | 122.2803 | 2 | Intertidal/ mangrove | 29/08/2018 | Green turtle observed |
| Roebuck Bay | 8 | -18.0321 | 122.2662 | 2 | Ocean | 29/08/2018 | Sea snake observed |
| Roebuck Bay | 9 | -18.0551 | 122.3700 | 1 | Intertidal | 30/08/2018 | Sea snake observed |
| Roebuck Bay | 10 | -18.0351 | 122.3245 | 2 | Ocean | 30/08/2018 | Flatback turtle observed |
| Roebuck Bay | 11 | -18.0084 | 122.2860 | 2 | Ocean | 30/08/2018 | Sea snake and flatback turtle observed |
| Roebuck Bay | 12 | -18.0077 | 122.2113 | 2 | Ocean | 30/08/2018 | |
| Cooktown | 1 | -15.3106 | 145.2906 | 5 | Creek/ mangrove | 13/02/2020 | Crocodiles have been previously observed here. |
| Cooktown | 2 | -15.3166 | 145.2857 | 5 | Creek/ wetland | 13/02/2020 | Crocodiles have been previously observed here. |

| | | | | | | | |
|---|---|---|---|---|---|---|---|
| Cooktown | 3 | -15.3184 | 145.2843 | 5 | Creek/ wetland | 14/02/2020 | Crocodiles have been previously observed here. |
| Cooktown | 4 | -15.3239 | 145.2829 | 5 | Creek/ mangrove | 14/02/2020 | Crocodiles have been previously observed here. |

### Table S2. eDNA taxa detection table

| Domain | Phylum | Class | Order | Family | Genus | Species | Roebuck Bay site # | Cooktown site # |
|---|---|---|---|---|---|---|---|---|
| Eukaryota | Arthropoda | Hexanauplia | Calanoida | Pontellidae | Labidocera | Labidocera japonica | 1 | |
| Eukaryota | Chordata | Actinopterygii | Atheriniformes | Atherinidae | Atherinomorus | | | 1 |
| Eukaryota | Chordata | Actinopterygii | Atheriniformes | Atherinidae | Atherinomorus | Atherinomorus duodecimalis | | 4 |
| Eukaryota | Chordata | Actinopterygii | Atheriniformes | Melanotaeniidae | Melanotaenia | | 8, 9 | 1, 3, 4 |
| Eukaryota | Chordata | Actinopterygii | Atheriniformes | Pseudomugilidae | Pseudomugil | Pseudomugil gertrudae | | 1, 2 |
| Eukaryota | Chordata | Actinopterygii | Aulopiformes | Synodontidae | Trachinocephalus | Trachinocephalus myops | | 4 |
| Eukaryota | Chordata | Actinopterygii | Beloniformes | Hemiramphidae | Hyporhamphus | Hyporhamphus quoyi | 4 | |
| Eukaryota | Chordata | Actinopterygii | Blenniiformes | Blenniidae | Omobranchus | Omobranchus elongatus | 4 | |
| Eukaryota | Chordata | Actinopterygii | Centrarchiformes | Terapontidae | Leiopotherapon | Leiopotherapon aheneus | 12 | 4 |
| Eukaryota | Chordata | Actinopterygii | Centrarchiformes | Terapontidae | Terapon | Terapon jarbua | | 1, 3, 4 |
| Eukaryota | Chordata | Actinopterygii | Clupeiformes | Chirocentridae | Chirocentrus | | 3, 5, 6, 9, 10 | |
| Eukaryota | Chordata | Actinopterygii | Clupeiformes | Chirocentridae | Chirocentrus | Chirocentrus dorab | 5 | |
| Eukaryota | Chordata | Actinopterygii | Clupeiformes | Clupeidae | Nematalosa | | 2, 3, 4, 5, 6, 9, 11, 12 | |
| Eukaryota | Chordata | Actinopterygii | Clupeiformes | Clupeidae | Sardinella | | 4, 5, 6, 7 | 4 |
| Eukaryota | Chordata | Actinopterygii | Clupeiformes | Engraulidae | Encrasicholina | | 1, 8 | 4 |
| Eukaryota | Chordata | Actinopterygii | Ephippiformes | Ephippidae | Platax | Platax orbicularis | 4 | |
| Eukaryota | Chordata | Actinopterygii | Gerreiformes | Gerreidae | Gerres | Gerres erythrourus | | 3, 4 |
| Eukaryota | Chordata | Actinopterygii | Gerreiformes | Gerreidae | Gerres | Gerres filamentosus | | 4 |
| Eukaryota | Chordata | Actinopterygii | Gobiiformes | Eleotridae | Eleotris | | | 1, 4 |
| Eukaryota | Chordata | Actinopterygii | Gobiiformes | Eleotridae | Hypseleotris | Hypseleotris compressa | | 1, 2, 3, 4 |
| Eukaryota | Chordata | Actinopterygii | Gobiiformes | Eleotridae | Ophiocara | Ophiocara porocephala | | 3, 4 |
| Eukaryota | Chordata | Actinopterygii | Gobiiformes | Eleotridae | | | | 1, 3, 4 |
| Eukaryota | Chordata | Actinopterygii | Gobiiformes | Gobiidae | Acentrogobius | Acentrogobius janthinopterus | | 4 |
| Eukaryota | Chordata | Actinopterygii | Gobiiformes | Gobiidae | Awaous | | | 1 |
| Eukaryota | Chordata | Actinopterygii | Gobiiformes | Gobiidae | Awaous | Awaous grammepomus | | 1 |
| Eukaryota | Chordata | Actinopterygii | Gobiiformes | Gobiidae | Cryptocentrus | Cryptocentrus cyanotaenia | 4 | |
| Eukaryota | Chordata | Actinopterygii | Gobiiformes | Gobiidae | Mugilogobius | Mugilogobius mertoni | | 4 |

| | | | | | | | | |
|---|---|---|---|---|---|---|---|---|
| Eukaryota | Chordata | Actinopterygii | Gobiiformes | Gobiidae | Mugilogobius | | | 4 |
| Eukaryota | Chordata | Actinopterygii | Gobiiformes | Gobiidae | Psammogobius | Psammogobius biocellatus | | 4 |
| Eukaryota | Chordata | Actinopterygii | Gobiiformes | Gobiidae | | | 6 | |
| Eukaryota | Chordata | Actinopterygii | Labriformes | Labridae | Stethojulis | Stethojulis trilineata | 1 | |
| Eukaryota | Chordata | Actinopterygii | Lutjaniformes | Haemulidae | Plectorhinchus | Plectorhinchus gibbosus | | 4 |
| Eukaryota | Chordata | Actinopterygii | Lutjaniformes | Haemulidae | Pomadasys | Pomadasys kaakan | 5, 6, 7, 9 | |
| Eukaryota | Chordata | Actinopterygii | Lutjaniformes | Haemulidae | | | 3 | |
| Eukaryota | Chordata | Actinopterygii | Lutjaniformes | Lutjanidae | Lutjanus | Lutjanus fulviflamma | | 4 |
| Eukaryota | Chordata | Actinopterygii | Lutjaniformes | Lutjanidae | Lutjanus | Lutjanus argentimaculatus | | 1, 4 |
| Eukaryota | Chordata | Actinopterygii | Lutjaniformes | Lutjanidae | Lutjanus | | | 4 |
| Eukaryota | Chordata | Actinopterygii | Lutjaniformes | Lutjanidae | Lutjanus | Lutjanus carponotatus | 4 | |
| Eukaryota | Chordata | Actinopterygii | Lutjaniformes | Lutjanidae | Lutjanus | Lutjanus fulvus | 4 | |
| Eukaryota | Chordata | Actinopterygii | Mugiliformes | Mugilidae | Ellochelon | Ellochelon vaigiensis | 4 | 1, 4 |
| Eukaryota | Chordata | Actinopterygii | Mugiliformes | Mugilidae | Planiliza | Planiliza subviridis | | 3, 4 |
| Eukaryota | Chordata | Actinopterygii | Perciformes | Serranidae | Epinephelus | Epinephelus lanceolatus | 6, 11 | |
| Eukaryota | Chordata | Actinopterygii | Spariformes | Nemipteridae | Scolopsis | Scolopsis monogramma | 1, 4, 12 | |
| Eukaryota | Chordata | Actinopterygii | Synbranchiformes | Synbranchidae | | | | 1, 4 |
| Eukaryota | Chordata | Actinopterygii | Syngnathiformes | Callionymidae | Callionymus | Callionymus russelli | 5, 6, 8 | |
| Eukaryota | Chordata | Actinopterygii | Syngnathiformes | Mullidae | Upeneus | Upeneus tragula | | 4 |
| Eukaryota | Chordata | Actinopterygii | Syngnathiformes | Pegasidae | Pegasus | Pegasus volitans | 3 | |
| Eukaryota | Chordata | Actinopterygii | Tetraodontiformes | Diodontidae | Diodon | | 1 | |
| Eukaryota | Chordata | Actinopterygii | Tetraodontiformes | Monacanthidae | Paramonacanthus | Paramonacanthus choirocephalus | | 4 |
| Eukaryota | Chordata | Actinopterygii | | Pomacanthidae | Chaetodontoplus | Chaetodontoplus duboulayi | 4 | |
| Eukaryota | Chordata | Actinopterygii | | Pseudochromidae | Pseudochromis | Pseudochromis elongatus | 1 | |
| Eukaryota | Chordata | Actinopterygii | | Sillaginidae | Sillago | Sillago asiatica | | 2, 4 |
| Eukaryota | Chordata | Actinopterygii | | Sillaginidae | Sillago | | | 4 |
| Eukaryota | Chordata | Actinopterygii | | Sillaginidae | Sillago | Sillago schomburgkii | 4, 12 | 3, 4 |
| Eukaryota | Chordata | Actinopterygii | | Sillaginidae | Sillago | Sillago aeolus | 4, 5 | |
| Eukaryota | Chordata | Amphibia | Anura | Bufonidae | Rhinella | Rhinella marina | | 1, 2, 3, 4 |
| Eukaryota | Chordata | Amphibia | Anura | Hylidae | Litoria | Litoria bicolor | | 1, 3 |
| Eukaryota | Chordata | Amphibia | Anura | Hylidae | Litoria | Litoria microbelos | | 2, 3 |
| Eukaryota | Chordata | Aves | Charadriiformes | Laridae | | | 4 | |
| Eukaryota | Chordata | Aves | Charadriiformes | Scolopacidae | Tringa | Tringa brevipes | 4 | |
| Eukaryota | Chordata | Aves | Charadriiformes | Scolopacidae | | | 10 | |

| | | | | | | | | |
|---|---|---|---|---|---|---|---|---|
| Eukaryota | Chordata | Aves | Columbiformes | Columbidae | Ducula | Ducula melanochroa | | 4 |
| Eukaryota | Chordata | Aves | Coraciiformes | Alcedinidae | | | | 4 |
| Eukaryota | Chordata | Aves | Galliformes | Phasianidae | Gallus | Gallus gallus | 3, 7, 12 | 3 |
| Eukaryota | Chordata | Aves | Passeriformes | Meliphagidae | Nesoptilotis | Nesoptilotis leucotis | | 4 |
| Eukaryota | Chordata | Aves | Passeriformes | Meliphagidae | | | | 3, 4 |
| Eukaryota | Chordata | Reptilia | Squamata | Colubridae | Dendrelaphis | Dendrelaphis calligaster | | 3 |
| Eukaryota | Chordata | Reptilia | Squamata | Homalopsidae | | | | 1 |
| Eukaryota | Chordata | Reptilia | Squamata | Scincidae | Carlia | Carlia storri | | 1 |
| Eukaryota | Chordata | Reptilia | Squamata | Scincidae | Carlia | Carlia longipes | | 1 |
| Eukaryota | Chordata | Reptilia | Squamata | Scincidae | Saproscincus | Saproscincus basiliscus | 12 | 2 |
| Eukaryota | Chordata | Mammalia | Artiodactyla | Bovidae | Bos | Bos taurus | 3, 8 | 1, 2 |
| Eukaryota | Chordata | Mammalia | Artiodactyla | Delphinidae | Stenella | Stenella attenuata | 4, 5 | |
| Eukaryota | Chordata | Mammalia | Artiodactyla | Delphinidae | | | 5, 6 | |
| Eukaryota | Chordata | Mammalia | Artiodactyla | Suidae | Sus | Sus scrofa | 4, 10 | 2, 3, 4 |
| Eukaryota | Chordata | Mammalia | Carnivora | Canidae | Canis | | 4 | 1 |
| Eukaryota | Chordata | Mammalia | Primates | Hominidae | Homo | Homo sapiens | 1, 2, 3, 4, 5, 6, 7, 8, 9, 11, 12 | 1, 2, 3, 4 |
| Eukaryota | Chordata | Reptilia | Testudines | Chelidae | Emydura | | | 4 |
| Eukaryota | Chordata | Reptilia | Testudines | Chelidae | Myuchelys | Myuchelys latisternum | | 1, 3, 4 |
| Eukaryota | Chordata | Reptilia | Testudines | Cheloniidae | Chelonia | Chelonia mydas | 4 | |
| Eukaryota | Chordata | Reptilia | Testudines | Cheloniidae | Natator | Natator depressus | 1, 3, 4, 5, 9, 11 | |
| Eukaryota | Echinodermata | Ophiuroidea | Ophiurida | Ophiactidae | Ophiactis | Ophiactis lymani | 10 | |
| Eukaryota | Mollusca | Bivalvia | Ostreoida | Ostreidae | Booneostrea | Booneostrea subucula | 12 | |
| Eukaryota | Mollusca | Bivalvia | Ostreoida | Ostreidae | Crassostrea | Crassostrea zhanjiangensis | 4, 5, 6,7, 10, 12 | |
| Eukaryota | Mollusca | Bivalvia | Ostreoida | Ostreidae | Dendostrea | | 3, 4 | |
| Eukaryota | Mollusca | Bivalvia | Ostreoida | Ostreidae | Ostrea | Ostrea algoensis | 6 | 4 |
| Eukaryota | Mollusca | Bivalvia | Ostreoida | Ostreidae | Planostrea | Planostrea pestigris | | 4 |
| Eukaryota | Mollusca | Bivalvia | Ostreoida | Ostreidae | | | 4, 6, 12 | |
| Eukaryota | Mollusca | Bivalvia | Pectinoida | Pectinidae | Mimachlamys | Mimachlamys australis | 2 | |
| Eukaryota | Mollusca | Bivalvia | Pectinoida | Spondylidae | Spondylus | Spondylus victoriae | 1, 2, 8 | |
| Eukaryota | Mollusca | Bivalvia | Pholadomyoida | Laternulidae | Laternula | Laternula creccina | 8 | |
| Eukaryota | Mollusca | Bivalvia | Pterioida | Pinnidae | Atrina | Atrina pectinata | 9, 11 | |
| Eukaryota | Mollusca | Bivalvia | Pterioida | Pteriidae | Electroma | Electroma physoides | 1 | |

| | | | | | | | | |
|---|---|---|---|---|---|---|---|---|
| Eukaryota | Mollusca | Gastropoda | Cephalaspidea | Bullidae | Bulla | Bulla quoyii | 4, 6 | |
| Eukaryota | Platyhelminthes | Catenulida | | Stenostomidae | Stenostomum | Stenostomum simplex | | 1, 3, 4 |
| Eukaryota | Platyhelminthes | Catenulida | | Stenostomidae | Stenostomum | Stenostomum sthenum | | 1, 3 |
